# iGEX: an interactive Gene EXpression exploration platform for the analysis of transcriptomic and proteomic datasets

**DOI:** 10.64898/2026.10.24.761566

**Authors:** Saurabh Saha, James Whelan, Monika W. Murcha

**Affiliations:** School of Molecular Sciences C ARC Centre of Excellence in Plant Energy Biology, The University of Western Australia, Perth, WA 6009, Australia; State Key Laboratory of Plant Environmental Resilience, College of Life Science, Zhejiang University, Hangzhou, Zhejiang 310058, P.R. China

## Abstract

High-throughput transcriptomic and proteomic studies have generated a rapidly expanding collection of publicly available datasets. However, extracting biological insight from these resources often requires researchers to repeatedly download, reformat, integrate and analyse individual files using specialised tools. This process is time-consuming and often requires coding expertise. Here, we present iGEX, an interactive exploration platform that can incorporate publicly available and user-generated transcriptomic and proteomic datasets into a single visual workspace to simplify the analysis of multiple transcripts and/or proteins across numerous datasets. iGEX enables researchers to select gene/protein targets and experiments and compare expression patterns across biological conditions and experiments. User-generated data can be examined alongside numerous public resources such as EMBL-EBI Expression Atlas, Bgee, and PaxDb to determine whether any observed patterns are evident across experiments, species or experimental contexts. Users can visualise and move directly between data tables, principal component analysis, expression and correlation heatmaps, hierarchical clustering and volcano plots, all of which are editable and downloadable.

## Introduction

The past two decades has seen a rapid expansion in the generation and public availability of transcriptomic and proteomic datasets. This growth has been enabled by advances in high-throughput RNA sequencing (RNA-seq) and mass spectrometry, including greater processing capacity, higher sensitivity and faster acquisition, alongside a substantial reduction in cost (1–3). The scale and diversity of datasets is illustrated by projects such as the Genotype-Tissue Expression (GTEx) v8 project, which analysed 15,201 RNA-seq samples collected across 49 human tissues from 838 individuals (4), and the Accelerating Medicines Partnership for Alzheimer’s Disease (AMP-AD) proteomics study, which analysed more than 2,000 human brain samples and 400 cerebrospinal fluid samples (5). In plants, large and diverse transcriptomic and proteomic datasets have been generated for *Arabidopsis thaliana* (Arabidopsis) (6,7) and crops such as wheat (8) and maize (9), as well as from plants grown aboard the International Space Station (10).

Considerable effort has been directed towards curating these datasets to make them accessible via platforms such as the EMBL-EBI Expression Atlas (11), Bgee (12) and PaxDb (13). These platforms curate experimental metadata, standardise annotations and process data from different studies into consistent formats. However, extracting biological insight across these resources often requires researchers to identify and download multiple individual datasets, interpret different file structures, match gene or protein identifiers between sources, and reformat and reintegrate data. These challenges become more complex when researchers want to combine and compare experiments from multiple resources or compare public datasets with their own experimental datasets, which can be time-consuming and shift attention from the biological question to the mechanics of data analysis.

Here, we present iGEX, an interactive gene expression exploration platform that brings publicly available transcriptomic and proteomic datasets together into a single desktop application. iGEX allows users to import, organise, integrate and screen datasets taken from Expression Atlas (https://www.ebi.ac.uk/gxa/experiments) (11), Bgee (https://www.bgee.org/download/processed-expression-values) (12) and PaxDb (https://pax-db.org/download) (13), as well as individual user-generated datasets. iGEX allows users to customise and visualise data for expression heatmaps, correlation heatmaps, principal component analysis (PCA) plots, and volcano plots across multiple experiments and targets of interest. In addition, users can curate and save their own library to organise genes, proteins and experiments into biologically relevant groups and workspaces. Users can also identify homologous genes/proteins across species and compare their expression/abundance profiles across multiple datasets stored within their iGEX library. Furthermore, all analyses are presented in a user-friendly, downloadable and editable format for figure generation.

## Methods

### Software design and implementation

iGEX was implemented in Python 3.10.6 (14). Numerical and tabular operations were performed with NumPy 1.26.1 (15) and pandas 2.3.3 (16). Apache Arrow/PyArrow 24.0.0 was used to store and process tabular datasets in the Parquet data format (https://arrow.apache.org/). Openpyxl 3.1.5 (https://pypi.org/project/openpyxl/) provided Excel workbook support for importing and exporting datasets through pandas. The graphical user interface (GUI) and interactive plot editors were constructed using PySide6 6.11.0 (https://pypi.org/project/PySide6/), which provides Python bindings for the Qt framework, a set of C++ libraries for GUI development (https://www.qt.io/development/qt-framework). Interface icons were obtained from the Lucide project (https://github.com/lucide-icons/lucide) and used to design the graphical interface and data point markers within the PCA and volcano plots.

Pillow 10.1.0 (https://pillow.readthedocs.io/en/stable/) was used to record metadata required for PNG exports. Playwright 1.60.0 (https://playwright.dev/python/) was used to load scalable vector graphics (SVG) outputs in a hidden Chromium browser and capture high-resolution PNG images, in order to ensure consistency between SVG and PNG exports. PDF and SVG files were exported directly from the Chromium browser. OpenAI Codex, using GPT-5.5 and GPT-5.6 Sol (OpenAI: https://chat.openai.com), was used as a coding assistant; all generated code was manually reviewed and verified.

### Data import and storage

iGEX supports the import of data from Expression Atlas (11), Bgee (12) and PaxDb (13), as well as user-generated custom CSV, TSV and Excel datasets. Import scripts that automatically recognise the data source type were added to standardise data into a common internal structure organised by species, data type and source (Fig. S1). For custom datasets, users must follow the specified iGEX import format (Fig. S2). All data imported into the software are stored within a single Library.iGEXdat archive. Within this archive, data are organised into tables by species and data type, including the data matrix, experiment metadata, source records, user-defined gene/protein target lists, annotated names and associated gene/experiment groups. This structure allows curated lists and classifications to be reused across compatible datasets. Archive contents were manually inspected using a custom file browser to verify the correct import, organisation and storage of data.

### Dataset screening and quality control

A Dataset Manager was implemented to rapidly screen experiments within the dataset loaded into the software. For each experiment, iGEX calculates the total transcript/protein abundance by summing the abundance values of all genes/proteins. It reports the number of genes/proteins that have an abundance value greater than zero. iGEX imports the processed abundance values distributed by Expression Atlas (11), Bgee (12) and PaxDb (13) and does not apply any further normalisation. Users importing custom datasets are required to normalise their data prior to import.

Each experiment’s total abundance and transcript/protein count are compared with the corresponding median values across all experiments in the selected dataset. An orange warning is assigned to experiments when the total abundance differs from the median by more than 10% or the detected transcript/protein count differs by more than 30%. A red warning is assigned when the corresponding differences exceed 25% or 50%, respectively. These warnings are recalculated whenever a dataset is loaded.

### Dataset integration and identifier mapping

An internal identifier mapping archive, Map.iGEXdat, was implemented to support gene/protein identifier lookup. The current archive contains mappings between Ensembl gene identifiers (17) and UniProt accessions (18) across the taxa listed in Table S1. These mappings allow users to query datasets using either Ensembl gene identifiers or UniProt accessions.

Transcriptomic and proteomic datasets from Bgee, PaxDb, Expression Atlas, and user-generated datasets can be integrated directly when they share the same gene or protein identifiers. Since PaxDb and Expression Atlas use different identifier systems, mappings between Ensembl protein identifiers and UniProt accessions are included for the five species currently available in Expression Atlas (Table S1). Log files are generated during dataset merging for user review.

### Principal component analysis

PCA was implemented using singular value decomposition (SVD) in iGEX PCA Plot Studio using previously described methods (19–21). Any blank, missing and non-numeric abundance values were converted to zero before analysis. For Gene PCA analysis, transcripts/proteins were treated as observations and experiments as variables. For Experiment PCA analysis, the same gene-by-experiment matrix was transposed so that experiments were treated as observations and transcripts/proteins as variables.

SVD was performed using NumPy 1.26.1 (15), and principal component scores, loadings and the proportion of variance explained by each component were calculated from the resulting singular vectors and singular values as described previously (21). Points displayed in the PCA plot were grouped by k-means clustering using NumPy (15,22).

### Expression and correlation heatmaps

Expression heatmaps were generated directly from the transcript/protein abundance values in the selected dataset, without any additional normalisation. Any blank, missing, and non-numeric values were represented as zero.

Correlation heatmaps in iGEX Heatmap Studio were calculated using Pearson’s product-moment correlation coefficient (23). Gene correlation matrices compares transcript/protein abundance profiles across experiments, whereas experiment correlation matrices compares experiments across transcripts/proteins. Calculations were performed using pandas 2.3.3 (16). Correlation coefficients that could not be calculated due to insufficient variation or an insufficient number of valid observations were represented as zero.

Heatmap colours were determined using specified minimum, midpoint and maximum values and their corresponding colours. Values outside the user-defined range were assigned the corresponding minimum or maximum colour. Linear, logarithmic, gamma, sinusoidal or exponential scaling functions were implemented for users to control the distribution of intermediate colour values across the gradient.

Hierarchical clustering of genes/proteins and experiments was implemented using previously described linkage methods (24,25) through SciPy 1.11.3 (26). Four linkage methods were implemented: single, complete and average linkage using correlation distance, and Ward linkage using Euclidean distance (24,27). When user-defined gene or experiment groups were present, members were clustered independently within each group to preserve the defined group structures.

Dendrogram branch length transformations were implemented in iGEX by adjusting the original clustering distances returned by SciPy 1.11.3 (26). This included the original distances (true distances); normalised distances, where values were converted to a 0-1 range; logarithmic or square-root scaling to compress large distances; equal-level cladogram spacing; and distance clipping, where distances were capped at the 95^th^ percentile to limit the visual influence of unusually long branches. These options change the dendrogram’s appearance without altering the clustering.

### Differential expression visualisation

The iGEX Differential Plot Studio was implemented to visualise differential expression as volcano plots using paired log_2_ fold-change and P-value estimates. These values are displayed without further normalisation or statistical recalculation. For each comparison, log_2_ fold change was plotted on the x-axis and statistical significance, calculated as −log_10_(P-value), was plotted on the y-axis. Entries lacking a valid pair of fold-change and P-value estimates are excluded. For plotting, P-values reported as zero in the source dataset were assigned a minimum value of 2.225 × 10⁻³⁰⁸.

Genes were classified as positively changed, negatively changed or not significant according to user-defined fold-change and P-value thresholds. To maintain viewport performance with large datasets, the number of background genes was limited to 500 by default, but this number can be increased by the user. Controls are also available to selectively display genes with the largest positive and negative fold changes, as well as tools to quickly identify experiments in which genes of interest show significant changes according to user-defined thresholds.

### Orthologue identification

iGEX Orthoscope was introduced as a feature within the Heatmap Studio and Differential Plot Studio to support the analysis of genes/proteins across species. Orthoscope queries Ensembl BioMart (28) for annotated gene orthologues and for comparisons across different Ensembl taxonomic divisions, for proteomic datasets, or in cases where BioMart does not support the selected species, Orthoscope retrieves protein sequences from UniProt and searches the target taxon using NCBI BLAST with default parameters (29,30).

Orthologue matches are presented alongside their E-values, query coverage and confidence scores within the target dataset. Hits are ranked by confidence, with the top hit for each query gene/protein automatically selected, while additional matches remain available for user review and selection.

## Results

### Overview of the iGEX interface and data analysis workflow

iGEX was developed as a single modular desktop application for the import, organisation, exploration and visualisation of multiple transcriptomic and proteomic datasets. The software is divided into four modules or studios: “Explorer” for dataset selection and curation, “PCA Plot Studio” for multivariate analysis, “Heatmap Studio” for expression and correlation heatmaps, and “Differential Plot Studio” for visualising differential expression datasets (Fig. 1). Together, these modules allow a user to move from dataset selection and curation to multivariate and gene expression analysis within a continuous workflow. Users can export publication-quality figures as high-resolution images and Excel spreadsheets containing the underlying data and analysis results.

**Figure 1.**
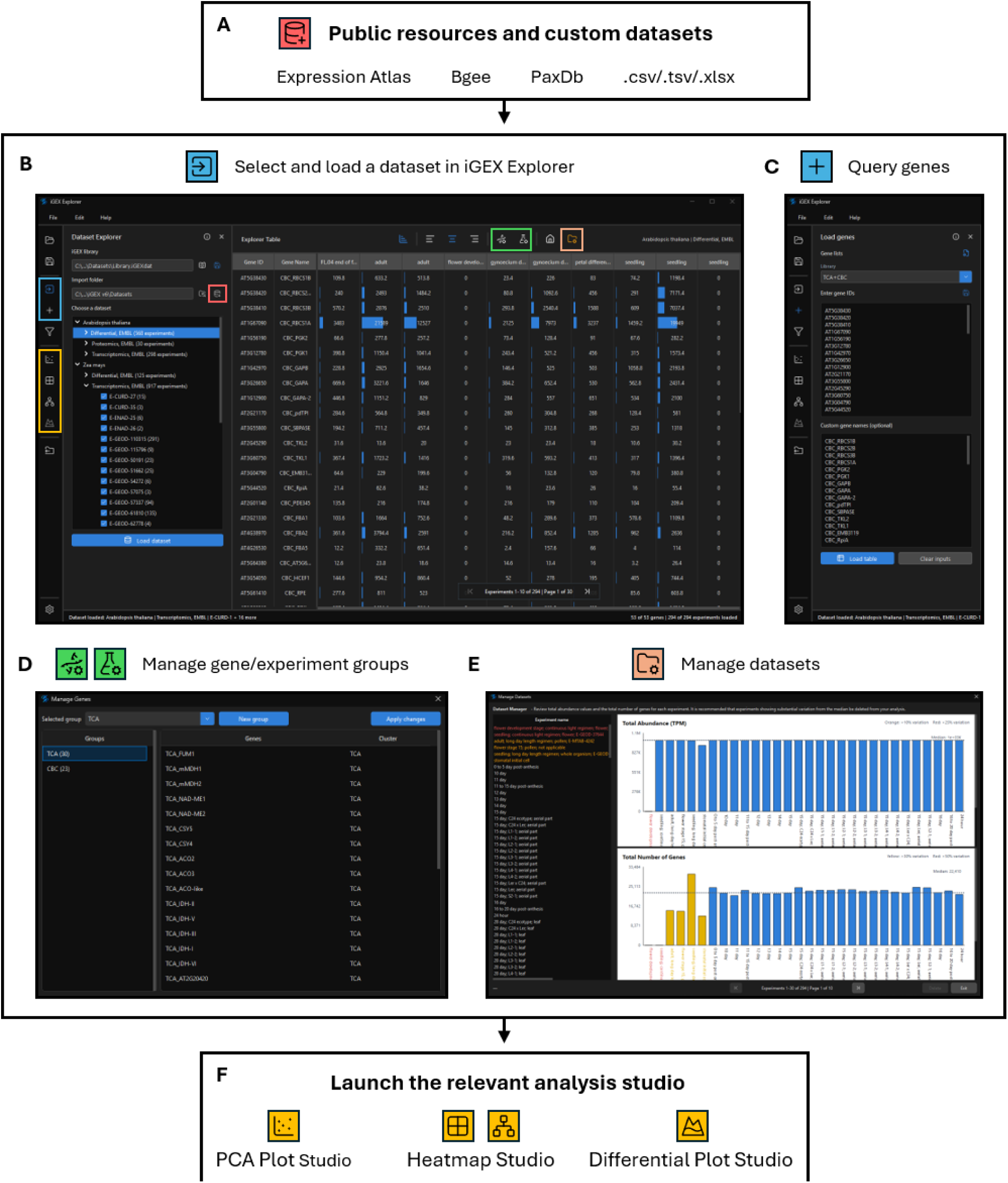
Overview of the iGEX interface and data analysis workflow. **(A)** Transcriptomic, proteomic and differential expression datasets from public resources or user-generated files can be imported into iGEX. **(B)** The iGEX Explorer interface serves as the central workspace for importing, loading and organising datasets. Queried genes/proteins are displayed as rows, while experiments within the selected dataset are displayed as columns. Datasets within the iGEX library are organised hierarchically by species, data type and source. **(C–E)** Interfaces for querying genes/proteins, managing gene and experiment groups, and screening datasets, respectively. **(F)** Selected data can be transferred to the PCA Plot Studio, Heatmap Studio or Differential Plot Studio. Coloured icons and outlines identify the corresponding controls within the main iGEX Explorer interface: red for importing datasets, blue for selecting datasets and querying genes/proteins, green for managing gene and experiment groups, orange for screening and managing datasets, and yellow for launching the corresponding analysis studios.

The iGEX Explorer serves as the central interface for selection of gene/protein targets and experimental datasets (Fig. 1). To begin, users must download datasets from public databases such as Expression Atlas (11), Bgee (12) and PaxDb (13), or prepare their own according to the specified iGEX format for custom datasets (Fig. 1A and S2). Once saved in a local folder, these datasets can be imported into the iGEX library, where iGEX identifies the data source, standardises the data structure and stores the datasets for subsequent analysis (Fig. S1). The Dataset Explorer icon in the left sidebar opens the dataset selection and import panel, where datasets within the iGEX library are organised by species, data type and source (Fig. 1B). The import dataset icon within this panel, outlined in red, is used to add new datasets to the library (Fig. 1B). After selecting a dataset, the Load Genes icon in the left sidebar can be used to open the gene/protein query panel (Fig. 1C), where users can enter compatible gene or protein identifiers and customise names.

Experiments are automatically assigned into groups using metadata tags provided by the original data source. For example, Expression Atlas supplies configuration files containing semicolon-separated experimental annotations such as “seedling” and “flowering stage” (11). These automatically assigned experiment groups can be individually reviewed and modified using Manage Experiments, while custom gene groups can be created using Manage Genes. The corresponding controls in the top toolbar are highlighted in green (Fig. 1B), with the Manage Genes interface shown as an example (Fig. 1D). Groups and individual genes or experiments can be renamed or deleted using the right-click menus.

Individual experiments loaded into iGEX Explorer can be manually screened via the Manage Datasets icon highlighted in orange (Fig. 1B, E). For each experiment, the iGEX interface displays the total abundance and number of detected genes/proteins and compares these values with the dataset medians. Abundance values represent measures such as transcripts per million (TPM) for transcriptomic data or intensity-based absolute quantification (iBAQ) for proteomic data. Experiments that deviate substantially from the median are highlighted and the Manage Datasets icon turns orange when flagged experiments are present (Fig. 1E). Users can retain, mark as reviewed or permanently delete experiments from their library using the right-click menu. The queried genes/proteins and selected experiments can be analysed in the relevant analysis studio using the controls highlighted in yellow (Fig. 1B, F).

### PCA Plot Studio: Multivariate exploration and screening of expression datasets

The PCA Plot Studio allows users to explore patterns and variation across genes/proteins and experiments within datasets loaded into iGEX Explorer (Fig. 2A). The controls highlighted in red in the top toolbar allow users to switch between two PCA modes: Gene PCA, which compares gene/protein expression profiles across selected experiments, or Experiment PCA, which compares experiments using the selected genes/proteins. The PCA Analysis panel, accessed through the icon highlighted in blue in the left sidebar, allows users to choose the number of principal components calculated and clusters displayed. Users can also restrict the analysis to groups previously defined in iGEX Explorer through the Gene group and Experiment group dropdown menus (Fig. 2A). In addition, “background” genes or experiments (a randomly chosen subset of genes/proteins or experiments from the source dataset) can be included in the PCA Plot analysis to determine whether any observed variation is specific to the selected data or reflects broader variation within the dataset (Fig. 2A).

**Figure 2.**
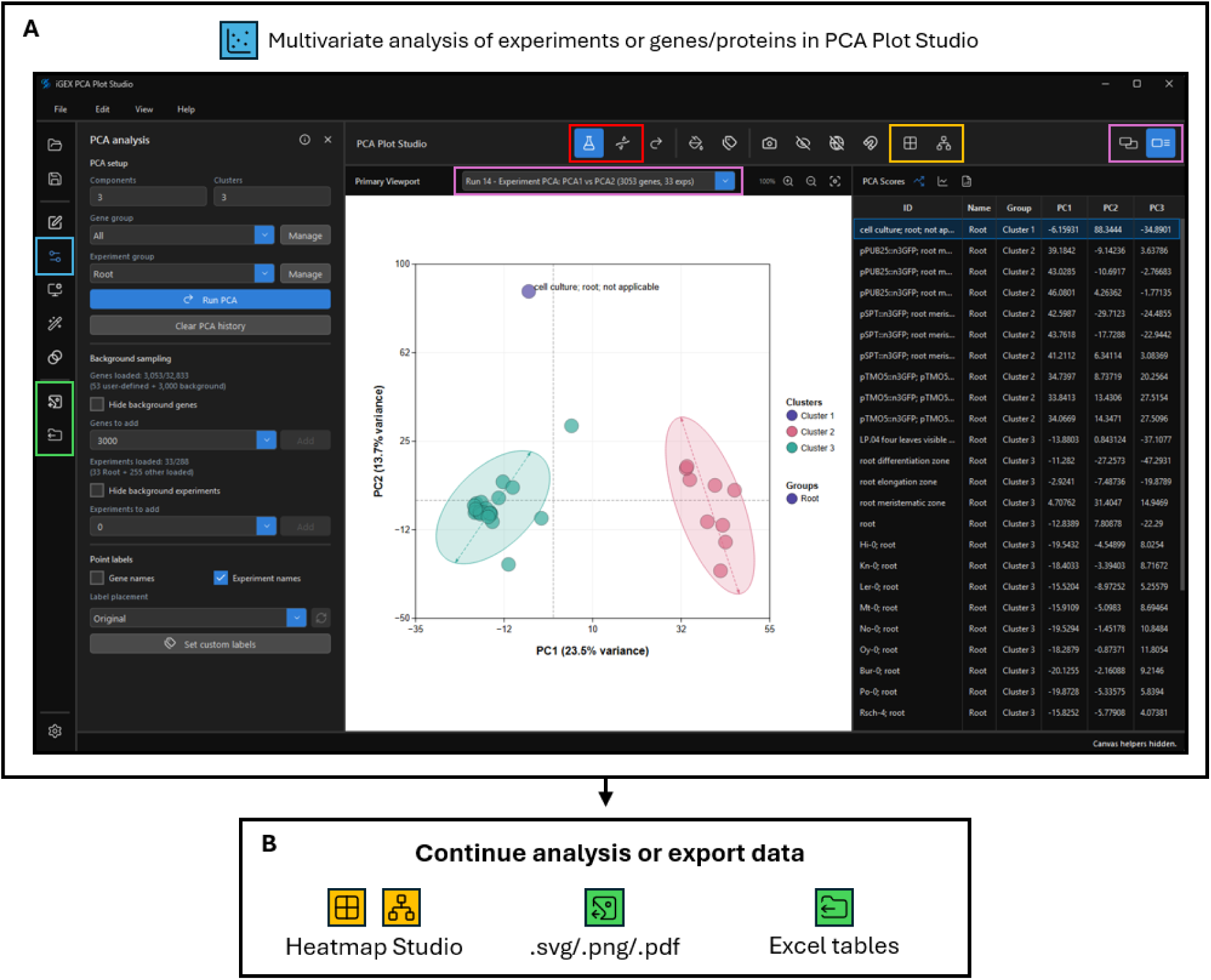
Overview of the iGEX PCA Plot Studio interface and analysis workflow. **(A)** Experiment PCA of 33 *Arabidopsis thaliana* experiments assigned to the “root” group based on metadata tags provided by Expression Atlas. The analysis used 53 user-selected genes and 3,000 additional background genes. Each point represents an experiment, and point colours indicate cluster assignments. The PCA Analysis panel and corresponding PCA scores table are shown alongside the PC1-PC2 plot. Shaded regions represent covariance ellipses calculated from the PC1 and PC2 scores within each cluster. **(B)** Curated gene or experiment groups can be transferred to Heatmap Studio for expression or correlation analysis. PCA plots can also be exported as figures, while the underlying data can be exported to Excel for further analysis. Coloured icons and outlines identify the corresponding controls within the PCA Plot Studio: blue for opening the PCA Analysis panel to adjust analysis settings, editing gene/experiment groups and adding background genes or experiments; red for switching between Gene PCA and Experiment PCA modes; purple for selecting previous PCA runs via the viewport dropdown menu and displaying plots side by side or alongside analysis tables; yellow for transferring curated data directly to the Heatmap Studio for expression or correlation analysis; and green for exporting plots as SVG, PNG or PDF files and analysis tables as Excel spreadsheets.

When the analysis settings are modified, the rerun icon turns orange to indicate that the PCA must be recalculated. Previous calculations are retained as separate PCA runs in the viewport dropdown menu, highlighted in purple, allowing users to revisit analyses performed with different gene sets, experiment selections or PCA settings. The view controls at the top right, also highlighted in purple, allow users to display two plots side by side for comparisons of PCA runs, or a single plot accompanied by PCA scores, loadings or explained-variance tables (Fig. 2A).

To illustrate a general Experiment PCA workflow, 33 Arabidopsis experiments were assigned to the “root” experiment group based on metadata tags provided by Expression Atlas. 53 user-selected genes and an additional 3,000 background genes were analysed (Fig. 2A). Each point represents an individual experiment. Most experiments formed two clusters, shown in pink and teal, which were separeted along PC1 (Fig. 2A). A single root cell-culture entry, shown in purple, was separeted along PC2 (Fig. 2A). This example illustrates how PCA can help users identify differences within a selected experiment group and review their experimental context before further analysis.

Following this review, individual genes or experiments can be excluded from the active analysis or moved to another group using the right-click menu on the corresponding point in the plot, without deleting data from the underlying library. Users can access the other sidebar menus to customise the plots and export them as high-resolution PNG images or editable vector graphics (SVG or PDF) (Fig. 2B). Users can directly transfer curated gene or experiment groups to Heatmap Studio for expression or correlation analysis using the icons indicated (Fig. 2A, B).

### Heatmap Studio: Expression, correlation and hierarchical clustering

The iGEX Heatmap Studio allows users to visualise transcript and protein abundance data as heatmaps and to compare expression profiles using correlation heatmaps (Fig. 3A). Expression heatmaps display the abundance values supplied by the source dataset, without further normalisation by iGEX. For correlation analysis, users can select either genes/proteins or experiments, with the resulting heatmap displaying their pairwise Pearson correlation coefficients (23).

**Figure 3.**
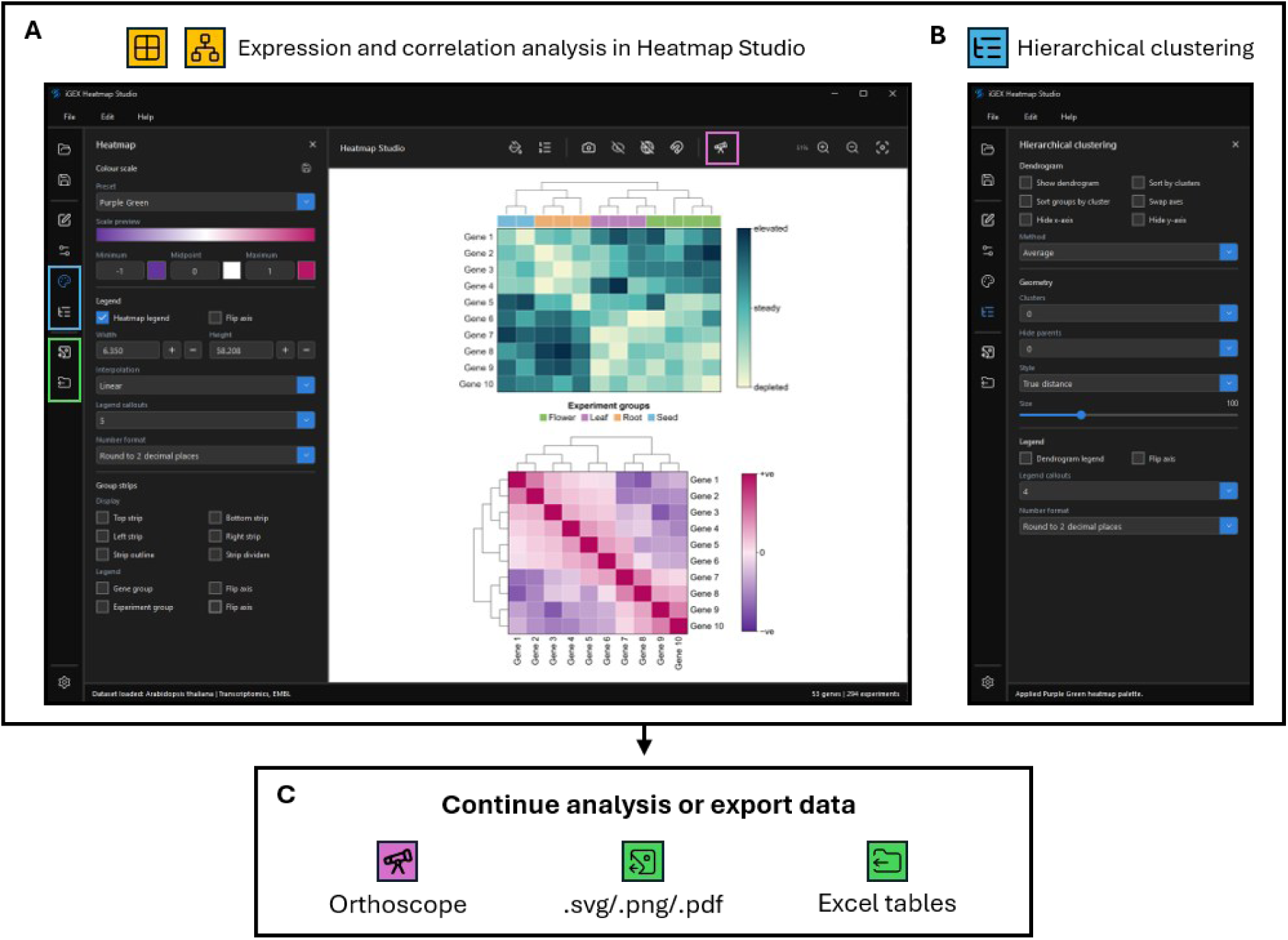
Expression, correlation and hierarchical clustering analysis in iGEX Heatmap Studio. **(A)** The Heatmap panel allows users to modify the displayed colours, select the method used to interpolate between colours, and adjust the number of callouts displayed on the heatmap legend. Examples of an expression heatmap (top) and a correlation heatmap (bottom) generated in iGEX are shown. **(B)** The Hierarchical Clustering panel provides controls for sorting genes/proteins or experiments, selecting the clustering method and adjusting the dendrogram display. (**C)** The iGEX Orthoscope interface can be accessed using the top toolbar icon to perform gene/protein identifier lookups across species with transcriptomic or proteomic datasets stored in the iGEX library. Alternatively, heatmaps and their underlying data can be exported. Coloured icons and outlines identify the corresponding analysis and interface controls: yellow for expression and correlation analysis; blue for opening the Heatmap and Hierarchical Clustering panels to adjust colours, legend labels, sorting and dendrogram settings; purple for accessing Orthoscope; and green for exporting heatmaps as SVG, PNG or PDF files and underlying data as Excel spreadsheets.

Heatmap colours are assigned using the minimum, midpoint, and maximum values according to the colours defined in the Heatmap colour panel (Fig. 3A). Users can select the interpolation method and adjust the number and formatting of callouts displayed on the heatmap legend by using the dropdown menus in the Heatmap panel, through the corresponding icon highlighted in blue in the left sidebar (Fig. 3A). Hierarchical clustering can be applied to sort genes/proteins or experiments according to their expression or correlation profiles (Fig. 3B). The Hierarchical Clustering panel, also accessed through an icon highlighted in blue, allows users to select the clustering method, define the number of clusters and adjust the dendrogram size and distance scaling (Fig. 3B). If gene or experiment groups have previously been defined, the clustering can be performed within the groups to preserve the group organisation (Fig. 3B).

To allow for analysis across species, users can access iGEX Orthoscope through the icon highlighted in purple in the top toolbar (Fig. 3C). If the iGEX library contains transcriptomic or proteomic datasets for more than one species, users can perform a gene/protein orthologue lookup against a target species using Ensembl BioMart (28) or NCBI BLAST (29,30). Ensembl BioMart returns previously annotated orthologues, whereas BLAST returns candidate homologues alongside E-values, query coverage and confidence scores. Users can select homologues to be analysed directly in a new iGEX Explorer page.

### Differential Plot Studio: Interactive exploration of differential expression datasets

To support the analysis of differential transcript expression datasets, the iGEX Differential Plot Studio was implemented to allow users to browse and visualise multiple differential datasets within a single interface (Fig. 4). Datasets can be downloaded from Expression Atlas (11) or prepared by users according to the specified iGEX format (Fig. S2). Each experimental comparison is presented as a volcano plot with log₂ fold-change on the x-axis and −log₁₀(P-value) on the y-axis (Fig. 4A). Users can open the Volcano Plot Settings panel using the icon highlighted in blue to adjust the P-value and log_2_ fold change thresholds as required (Fig. 4A). These thresholds automatically reclassify genes as positively changed, negatively changed or not significant.

**Figure 4.**
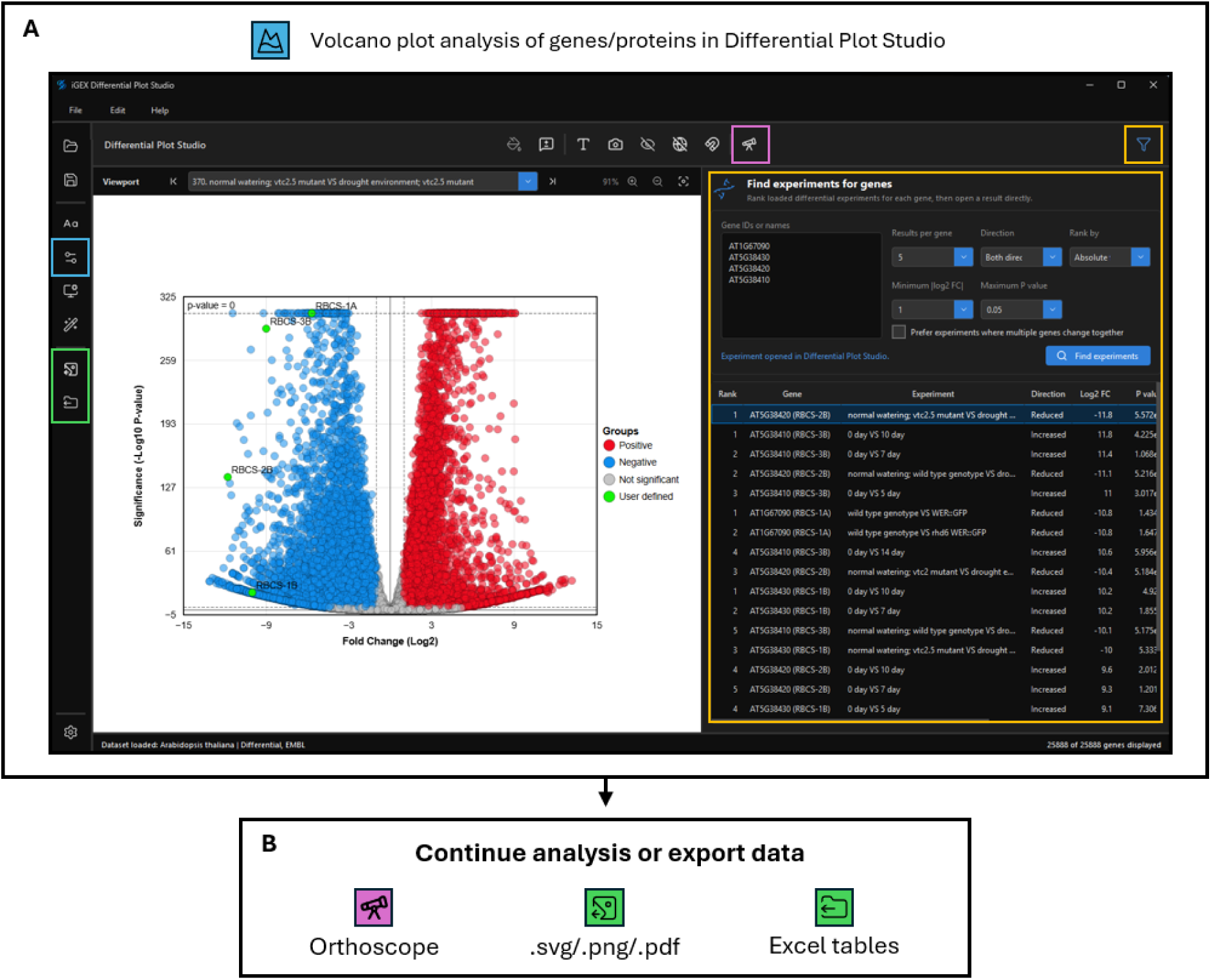
Screening differential expression datasets in the iGEX Differential Plot Studio. **(A)** Four *Arabidopsis thaliana* RBCS genes were queried in the Filter Experiments panel. Up to five differential comparisons per gene were retrieved using |log₂ fold change| ≥ 1 and P ≤ 0.05. The volcano plot compares dry seed transcriptomic expression against seeds imbibed for 48 h at 4°C in the ascorbate-deficient *vtc2*-4 mutant from E-MTAB-5103 dataset (31). Red and blue points indicate significantly increased and reduced expression, respectively; grey indicates genes below the significance thresholds (|log₂ fold change| > 1; P < 0.05); and green highlights the four queried RBCS genes. The upper dashed line marks points reported with a P-value of zero in the source dataset, which were assigned a minimum P-value of 2.225 × 10⁻³⁰⁸. **(B)** The iGEX Orthoscope interface can be accessed using the top toolbar icon to perform gene/protein identifier lookups across species with differential datasets stored in the iGEX library. Alternatively, volcano plots and their underlying data can be exported. Coloured icons and outlines identify the corresponding controls: blue for opening the layout panel, yellow for finding experimental comparisons, purple for Orthoscope and green for export.

To identify experimental conditions associated with significant changes in the abundance of genes of interest, users can open the Find experiments for genes panel through the filter icon highlighted in yellow in the top RHS toolbar (Fig. 4A). Users can enter query genes using their respective accession IDs, and specify a minimum absolute log_2_ fold-change and maximum P-value thresholds to identify experiments in which these genes show significant changes (Fig. 4A). Matching differential comparisons can be sorted by any of the table columns, and selecting a result row will automatically open the corresponding differential comparison. In the example shown (Fig. 4), four Arabidopsis genes encoding the small subunits of Rubisco, *RBCS1A*, *RBCS1B*, *RBCS2B* and *RBCS3B*, were used as queries against the preloaded differential datasets using thresholds of |log₂ fold change| ≥ 1 and P ≤ 0.05 and ranked by absolute fold change. The highest-ranked comparison is shown in the volcano plot, with the four queried genes highlighted in green (Fig. 4A). The selected experiment compared dry seeds with seeds imbibed for 48 h at 4 ° C in the ascorbate-deficient Arabidopsis *vtc2*-4 mutant (31). Negative log₂ fold changes indicate lower transcript abundance in dry seeds relative to imbibed seeds. Using this method we identified significant increases in *RBCS* transcript abundance in a separate study comparing dark-grown Arabidopsis cell cultures at day 0 with cultures sampled 5, 7, 10 and 14 days after exposure to light to induce chloroplast development (32) showing positive changes up to approximately 3600-fold (log_2_ fold change = 11.8) in the results table highlighted in yellow (Fig. 4A).

The iGEX Orthoscope module was implemented in the Differential Plot Studio to allow users to perform gene orthologue lookups across species and can be accessed through the icon highlighted in purple in the top toolbar (Fig. 4B).

### An example workflow: Conserved expression patterns of energy metabolism in plants

To demonstrate an integrated user case for iGEX, we examined transcript expression for numerous genes associated with the Calvin–Benson cycle (CBC) and tricarboxylic acid (TCA) cycle in Arabidopsis and *Zea mays* (maize) (33–36) across publicly available RNA-seq experiments spanning different tissues, developmental stages and experimental conditions. Arabidopsis CBC and TCA cycle gene identifiers were downloaded from AraCyc through the Plant Metabolic Network (37,38). A total of 82 pathway associated gene identifiers were obtained and filtered for experimental subcellular localisation data using SUBA5 (39). As the AraCyc pathway annotations include several paralogous isoenzymes that function in different compartments, genes encoding plastid-localised CBC or mitochondrion-localised TCA cycle proteins were prioritised, resulting in a final list of 53 genes (Table S2).

Transcriptomic RNA-seq data from 295 Arabidopsis experiments and 905 maize experiments were downloaded from Expression Atlas (11) and imported into iGEX Explorer. Seven Arabidopsis and 28 maize experiments were flagged by iGEX for substantial deviations in transcript abundance/total number of genes and were removed (Fig. S3). The Arabidopsis experiments were prefiltered using metadata tags and grouped manually into 10 distinct groups based on shared tags such as whole plant, leaf, root, seedling, etc. PCA was performed on the Arabidopsis experiments using the 53 query genes with 5,000 randomly selected background genes (Fig. 5A). The PCA plot showed broad variation amongst the experiments, with groups tagged as root, leaf, flower/reproductive, embryo/endosperm and genotype/line forming distinct clusters (Fig. 5A, arrows). However, some experiments deviated from their respective clusters, suggesting that the automated assignment of experimental groups based on metadata tags only provides a starting point, and users should verify the experimental conditions to manually curate these groups as required.

**Figure 5.**
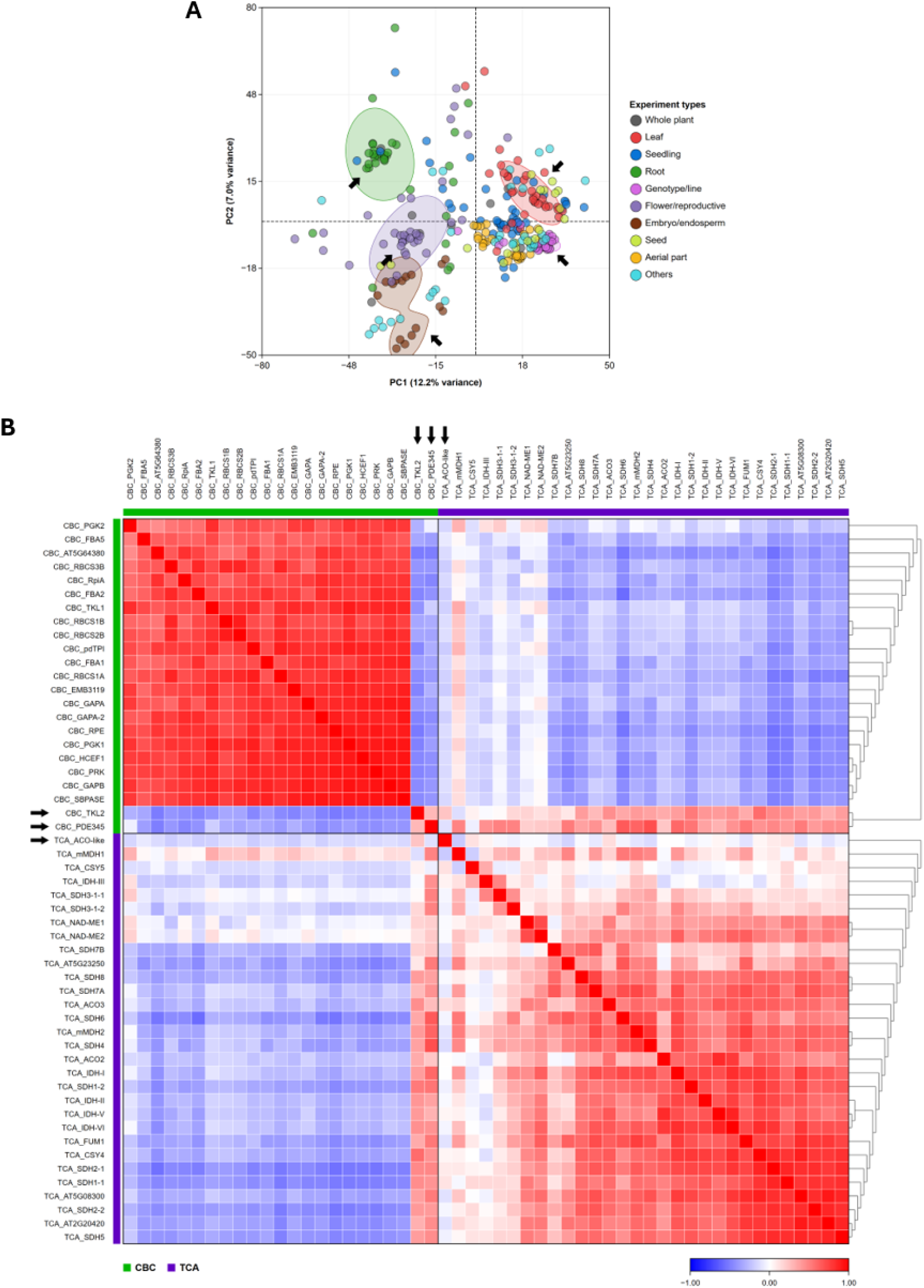
Expression patterns of Calvin–Benson cycle (CBC) and tricarboxylic acid (TCA) cycle genes in *Arabidopsis thaliana.* **(A)** PCA plot of 288 transcriptomic experiments imported from Expression Atlas (11). The analysis included a total of 53 CBC and TCA cycle genes, along with 5,000 randomly selected background genes. Experiments were automatically prefiltered using the metadata tags and manually grouped by common tags in iGEX. Shaded regions mark areas of highest point density within the corresponding experiment groups, and arrows highlight selected regions. **(B)** Pearson correlation matrix of the 53 CBC and TCA cycle genes across the 288 experiments. The genes were grouped into CBC and TCA and then ordered within each group by single-linkage hierarchical clustering. Green and purple annotation strips identify CBC and TCA genes, respectively. Dendrograms are displayed as equal-level cladograms. Red indicates positive correlation, white indicates no correlation and blue indicates negative correlation (−1 to 1).

A Pearson correlation matrix was calculated for the 53 query genes across the 288 experiments, and the genes were ordered within the user-defined CBC and TCA cycle groups using single-linkage hierarchical clustering (Fig. 5B). Most CBC genes formed a strongly positively correlated module on the top LHS, as did most TCA cycle genes (bottom RHS). However, several exceptions were observed. The genes encoding the plastidial transketolase TKL2 (AT2G45290) and the plastidial fructose-bisphosphate aldolase PDE345 (AT2G01140; also known as FBA3) negatively correlated with most CBC genes (Fig. 5B, arrows), consistent with their tissue-specific specialisations as *TKL2* was observed to be more highly expressed in flowers, roots and stems compared to leaves (40), unlike *TKL1* which is the predominant transketolase expressed in photosynthetic leaf tissues (41). Similarly, *PDE345* was observed to be expressed predominantly in heterotrophic tissues, particularly vascular tissues in leaves and roots, with little expression in the leaf mesophyll, consistent with its role in plastidial glycolytic metabolism in roots rather than photosynthetic carbon fixation (42). Within the TCA cycle group, the genes encoding ACO2 and ACO3, which catalyse the conversion of citrate to isocitrate in the TCA cycle, were strongly positively correlated within the main TCA cycle module as expected (43). The putative aconitase-like (ACO-like) gene AT5G54950 was also observed to deviate from the main correlation pattern (Fig. 5B, arrow). This gene encodes for a short, 74 aa protein with a yet uncharacterised function and is unlikely to function in a manner similar to the canonical ∼900 aa aconitases, such as ACO2 (AT4G26970) or ACO3 (AT2G05710) (Table S2) (43). These examples highlight the potential of iGEX expression/correlation heatmaps to assist in identifying gene candidates for further experimental investigation.

The 53 Arabidopsis genes were submitted to iGEX Orthoscope to identify homologous genes in maize. Candidate homologues were obtained through Ensembl BioMart (28), with up to five candidates identified for each gene (Table S3). Matches with less than 50% query coverage were excluded, resulting in a final set of 60 maize CBC and TCA cycle gene candidates. We performed a similar PCA on 877 RNA-seq experiments in maize using these candidates along with 5,000 randomly selected background genes (Fig. 6A) and grouped the experiments using the same metadata-based workflow applied to the Arabidopsis dataset. Here, the PCA revealed broad variation among the maize experiments, whereas experiments within the whole plant, genotype/line, temperature, stem and “others” groups formed distinct clusters (Fig. 6A, arrows).

**Figure 6.**
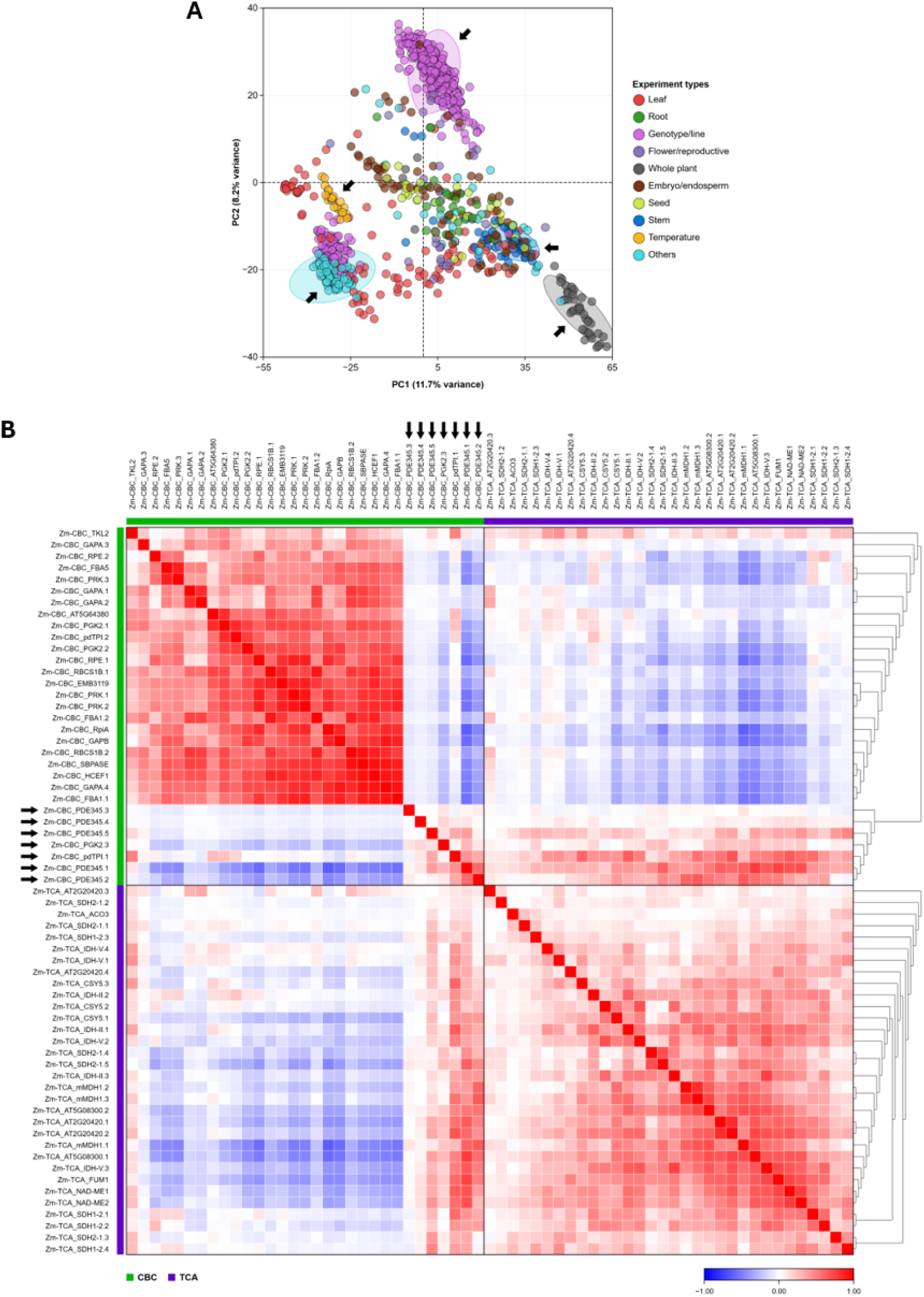
Expression patterns of Calvin–Benson cycle (CBC) and tricarboxylic acid (TCA) cycle homologues in *Zea mays*. **(A)** PCA plot of 877 transcriptomic experiments imported from Expression Atlas (11). The analysis included 60 maize homologues identified from the 53 Arabidopsis genes using iGEX Orthoscope, along with 5,000 randomly selected background genes. Experiments were automatically prefiltered using metadata tags and manually grouped by common tags in iGEX. Shaded regions mark areas of highest point density within the corresponding experiment groups, and arrows highlight selected regions. **(B)** Pearson correlation matrix of the 60 maize genes across the 877 experiments. Genes were grouped into CBC and TCA and ordered within each group by single-linkage hierarchical clustering. Green and purple annotation strips identify CBC and TCA genes, respectively. Dendrograms are displayed as equal-level cladograms. Red indicates positive correlation, white indicates no correlation and blue indicates negative correlation (−1 to 1).

A Pearson correlation matrix was calculated (Fig. 6B) and as previously observed for Arabidopsis, most CBC homologues formed a strongly positively correlated module at the top LHS, as did most TCA cycle homologues at the bottom RHS (Fig. 6B). Interestingly, five CBC homologues for PDE345 were negatively correlated with the main CBC module (Fig. 6B, arrows). This correlation was also seen within the Arabidopsis dataset (Fig. 5B, arrow). Cross-species comparisons identified similar patterns for homologues of the plastidial triose-phosphate isomerase (*pdTPI)*, which interconverts glyceraldehyde 3-phosphate and dihydroxyacetone phosphate (44), and the plastidial phosphoglycerate kinase 2 *(PGK2*), which catalyses the ATP dependent conversion of 3-phosphoglycerate to 1,3-bisphosphoglycerate during the CBC (Fig. 6B, arrows). Together, these observations demonstrate how biologically relevant expression patterns could be identified quickly across numerous genes and species using iGEX.

## Discussion

iGEX was developed for the exploration and visualisation of processed expression datasets, rather than raw-data processing. Many tools are already available for this, such as FragPipe, which provides workflows for peptide identification and protein quantification (45), and Salmon and RNAlysis that estimate transcript abundance from RNA-seq reads (46,47). iGEX simply allows users to import processed abundance matrices or precomputed fold-change and P-value tables in the supported iGEX format after the required quality control and normalisation. The advantage of iGEX is that it allows users to quickly determine whether experiments assigned to similar tissues or conditions exhibit comparable expression profiles using the PCA Plot Studio. Users can investigate if their own dataset or publicly available datasets deviate from other datasets obtained under comparable biological and experimental conditions. Conversely, users can search for specific experimental conditions associated with large changes in the expression of their genes of interest. Experiments can then be retained or excluded from the active analysis without modifying the underlying dataset. The iGEX Heatmap Studio allows for a quick and direct way to examine gene/protein abundance profiles across many experiments. Whilst datasets imported from Expression Atlas, Bgee and PaxDb contain normalised abundance values (11–13), this does not ensure that every experiment is suitable for comparison. For example, in one dataset, E-GEOD-37644 (48), only nine genes were observed with detectable transcript abundance values, yet their combined abundance was normalised to one million TPM, a value expected for the whole transcriptome (Fig. S3). As a result, the skewed values of a few select genes dominate the heatmap and mask true variation across the other experiments. Although this represents an extreme example, some experiments may still distort the heatmap scale and affect its interpretation and thus should be analysed with caution.

iGEX correlation heatmaps provide a rapid way to compare co-expression relationships among numerous gene sets and across multiple experiments. A strong correlation may indicate linked regulation or biological associations, although correlation alone does not demonstrate a direct functional relationship (49). It may provide insight into the roles of yet uncharacterised genes/proteins. Users should select biologically relevant experiment groups when comparing correlations between their genes of interest, as genes with similar tissue/developmental specific regulation will likely show strong correlations. Nevertheless, this analysis provides a useful starting point for generating experimental hypotheses. The Differential Plot Studio was developed to allow for the screening of multiple differential expression datasets quickly. Users can import their own differential data for analysis or explore the large collection of differential datasets available through Expression Atlas (11). As of the October 2025 release, Expression Atlas contained 14,587 differential comparisons across 51 species. These comparisons allow genes of interest to be visualised and ranked according to consistent P-value and log₂ fold-change thresholds, together with customised colours and callouts. This enables users to rapidly identify experimental conditions and/or mutant genotypes in which their genes are differentially expressed. The Orthoscope feature further extends this data analysis and allows users to quickly examine similar expression patterns across species.

The increasing availability and magnitude of transcriptomic and proteomic datasets have created challenges for their organisation, analysis and interpretation. Online resources such as Expression Atlas, Bgee and PaxDb provide invaluable access to curated experimental datasets, but they do not provide a single local platform for organising their resources alongside user-generated datasets (11–13). Moreover, long-term access to web-based tools and resources cannot be guaranteed, as many online services become redundant or unmaintained over time (50). Researchers commonly assemble datasets, prepare gene lists, develop scripts and rely on a range of available tools to examine their data. However, these workflows can become difficult to reproduce, especially after extended periods of time. For example, GENEVESTIGATOR enabled researchers to explore large collections of curated expression data, while PLEXdb allowed users to integrate datasets from multiple plant species, create and manage gene lists, and identify homologous genes between species (51,52). However, the discontinuation or changes in the availability of these resources, along with the requirement for paid licences for other tools such as Qlucore Omics Explorer (http://qlucore.com), has created a need for freely available software that can integrate these functions within a single workflow. We developed iGEX to address this need through a free interactive desktop environment that integrates publicly available datasets alongside user-generated data. iGEX is free to download and use and does not require registration or subscription. Users can organise and store datasets by species and data type within their own local libraries, identify homologous genes between species, and move directly between species and experimental groups for comparative analysis. iGEX provides a structured platform and workflow that supports incremental and reproducible exploration across publicly available and user-generated datasets.

## Conclusion and future directions

We developed iGEX as an interactive desktop environment that brings together the organisation, screening, analysis and visualisation of gene expression datasets. The dataset library was structured to organise data by type and species and to store user-defined gene sets and experiment groups locally. This allows datasets and analysis configurations to be reused while reducing reliance on online resources that may change or become unavailable. This initial release provides accessible workflows for dataset import, storage and screening, as well as PCA, expression, correlation and differential expression analyses, along with the identification of corresponding gene orthologues for cross-species comparisons.

We intend to continue expanding iGEX by implementing additional analysis methods, including multivariate analysis using uniform manifold approximation and projection (UMAP) (53), and extending support for single-cell expression data. As public databases expand and additional datasets become available, we will also extend the identifier-mapping library and develop import support for additional databases and resources. Ultimately, we intend for iGEX to serve as a comprehensive workbench for exploring and interpreting gene expression data.

## Data availability

The iGEX software is freely available for download from the iGEX website and GitHub (https://oneclickomics.github.io/iGEX/; https://github.com/Oneclickomics/iGEX).

Documentation describing the scientific methods and their implementation is available on the iGEX website. The data supporting the findings of this study are provided in the Supplementary Data accompanying the manuscript.

## Acknowledgements

The authors thank Benjamin Hearn-Thomas, Valencia Marissa, Yanqiao Zhu and Yang Zhao for providing valuable feedback and reporting critical software issues.

## Author Contributions

S.S, J.W and M.W.M conceived the project. S.S developed the software. All authors contributed to writing, reviewing and editing the manuscript, as well as testing and evaluating the software.

