## Supplementary Figures and Tables for "iGEX: an interactive Gene EXpression exploration platform for the analysis of transcriptomic and proteomic datasets"

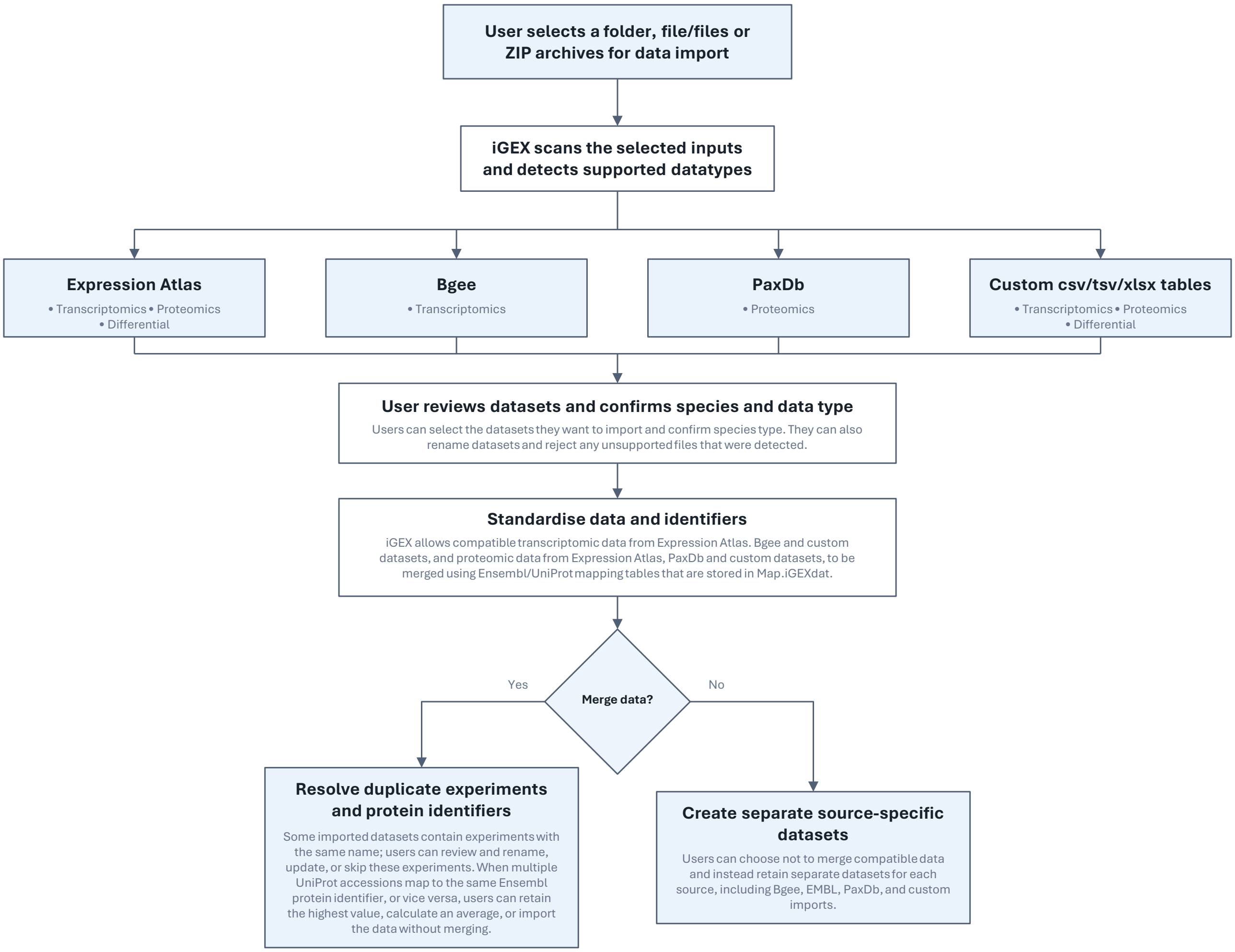


**Figure S1.** Data import, standardisation and storage workflow in iGEX. Users select one or more files, folders, or ZIP archives. iGEX identifies supported Expression Atlas, Bgee, PaxDb, and custom datasets, after which users review the detected datasets and confirm their species and data type (transcriptomic, proteomic, or differential). Compatible datasets can then be merged within the same species and data type or retained as source-specific datasets. When compatible proteomic datasets are merged, mappings stored in Map.iGEXdat are used to reconcile accession identifiers. If multiple abundance values resolve to the same target identifier, users can retain the highest value, calculate the mean, or import the records separately. The validated data are stored in Library.iGEXdat.

**A**

| Gene ID | Gene Name | Exp 1 | Exp 2 | Exp 3 | Exp 4 |
|---|---|---|---|---|---|
| AT1G01010 | LHY | 55 | 122 | 3.4 | 0 |
| AT2G21660 | GRF7 | 12 | 4 | 14.4 | 0 |
| AT3G18780 | ACT2 | 16 | 12.2 | 11.8 | 140 |
| AT5G52310 | RD29A | 0 | 14 | 23.5 | 105.4 |

**B**

| Gene ID | Gene Name | Exp 1.p-value | Exp 1.log2fold | Exp 2.p-value | Exp 2.log2fold |
|---|---|---|---|---|---|
| AT1G01010 | LHY | 0.86 | 0.2 | 0.91 | 0.7 |
| AT2G21660 | GRF7 | 0.65 | 0.5 | 0.96 | -0.2 |
| AT3G18780 | ACT2 | 0.85 | 0.3 | 0.92 | -0.5 |
| AT5G52310 | RD29A | 0.78 | -0.8 | 0.95 | -0.4 |

**Figure S2.** Required formats for custom datasets imported into iGEX. (A) Format for transcriptomic or proteomic measurements. The first two columns contain the gene or protein identifier and corresponding name, and each subsequent column contains numerical expression or abundance values for one experiment or measurement. (B) Format for differential data. The first two columns contain gene identifiers and names, followed by adjacent pairs of P-value and log2-fold-change columns for each comparison. Column names within each pair must share the same experiment label. Values are illustrative.

A

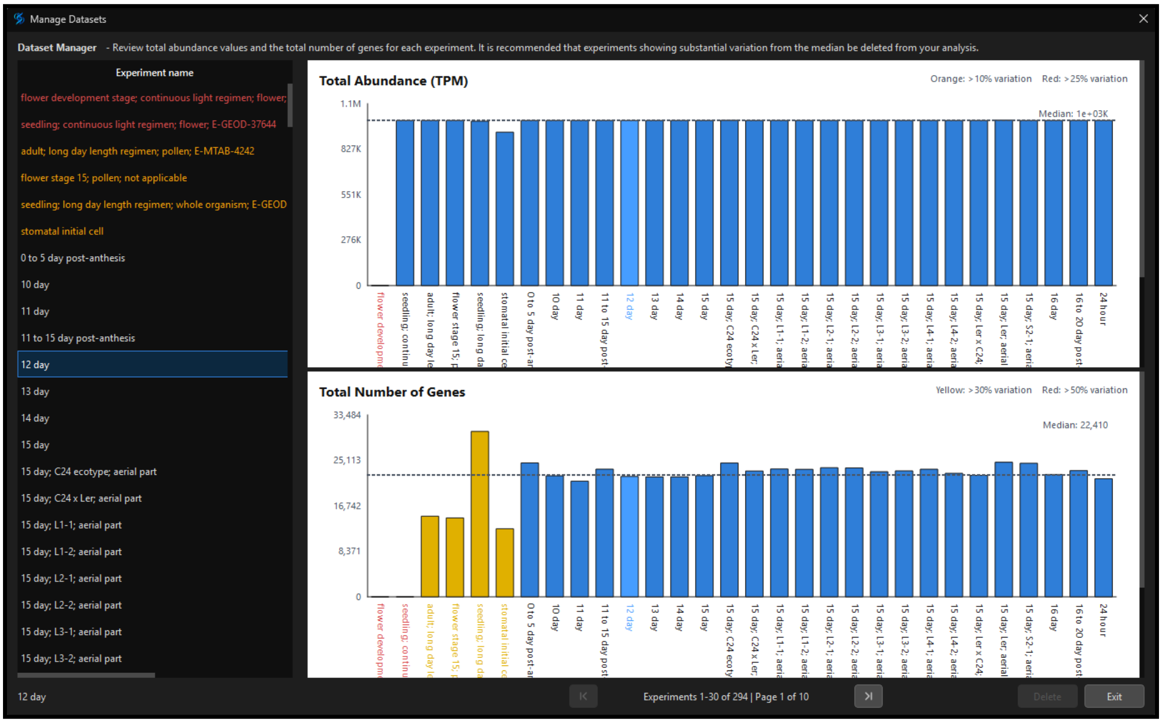


B

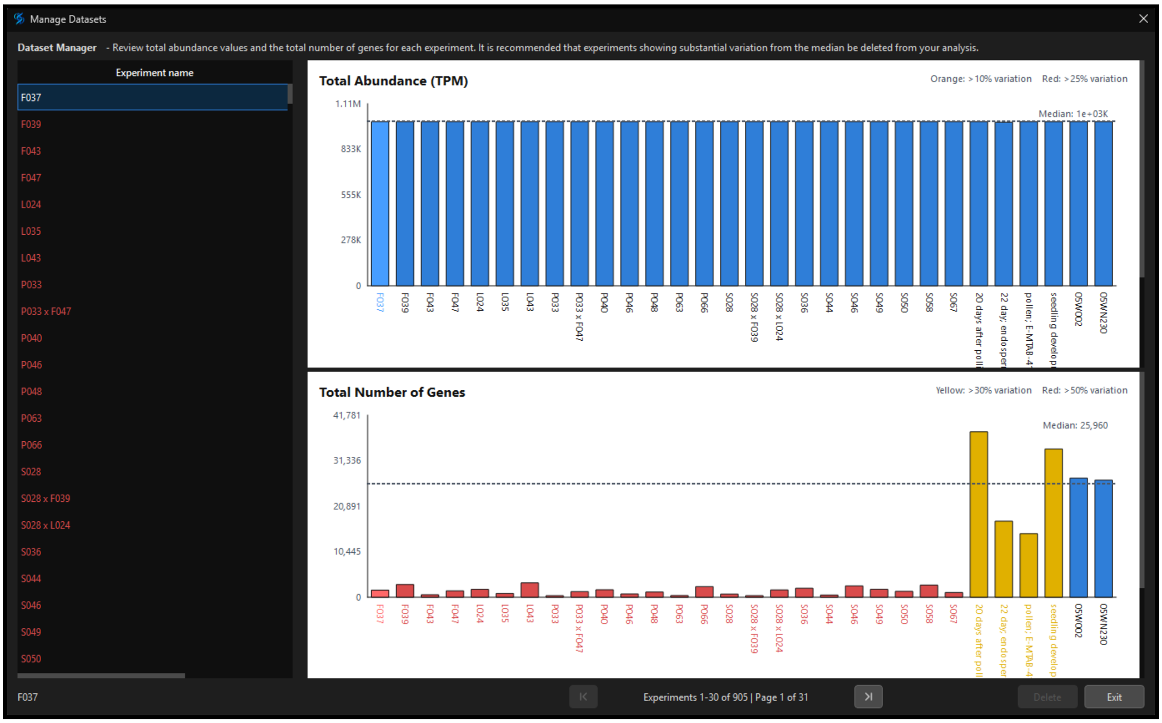


**Figure S3.** Screening RNA-seq datasets using the iGEX Manage Datasets interface. (A) *Arabidopsis thaliana* and (B) *Zea mays* datasets imported from Expression Atlas. The upper plots show total transcript abundance in transcripts per million (TPM), and the lower plots show the number of detected genes (TPM > 0) for each experiment. Dashed horizontal lines indicate the corresponding median across all experiments within each species. For total abundance, orange and red warnings indicate deviations from the median greater than 10% and 25%, respectively. For detected gene counts, yellow and red bars indicate deviations greater than 30% and 50%, respectively. Blue bars indicate experiments within deviation thresholds. Seven of 295 Arabidopsis experiments and 28 of 905 maize experiments were flagged and subsequently excluded from further analysis, leaving 288 and 877 experiments, respectively.

**Table S1.** Species and identifier types represented in Map.iGEXdat. G = Ensembl gene ID; U = UniProt accession; P = Ensembl protein ID. Blue rows identify the five species with proteomic datasets in Expression Atlas.

| Species | NCBI taxonomy ID | Ensembl division | BioMart dataset | Identifier types |
|---|---|---|---|---|
| *Aegilops tauschii* | 37682 | plants | atauschii_eg_gene | G; U |
| *Anolis carolinensis* | 28377 | ensembl | acarolinensis_gene_ensembl | G; U |
| *Arabidopsis lyrata* | 59689 | plants | alyrata_eg_gene | G; U |
| *Arabidopsis thaliana* | 3702 | plants | athaliana_eg_gene | G; U; P |
| *Beta vulgaris subsp. vulgaris* | 3555 | plants | bvulgaris_eg_gene | G; U |
| *Bos taurus* | 9913 | ensembl | btaurus_gene_ensembl | G; U |
| *Brachypodium distachyon* | 15368 | plants | bdistachyon_eg_gene | G; U |
| *Brassica napus* | 3708 | plants | bnapus_eg_gene | G; U |
| *Brassica oleracea* | 3712 | plants | boleracea_eg_gene | G; U |
| *Brassica rapa* | 3711 | plants | brapa_eg_gene | G; U |
| *Brassica rapa subsp. rapa* | 3711 | plants | brapa_eg_gene | G; U |
| *Caenorhabditis elegans* | 6239 | ensembl | celegans_gene_ensembl | G; U |
| *Chlamydomonas reinhardtii* | 3055 | plants | creinhardtii_eg_gene | G; U |
| *Chlorocebus sabaeus* | 60711 | ensembl | csabaeus_gene_ensembl | G; U |
| *Danio rerio* | 7955 | ensembl | drerio_gene_ensembl | G; U |
| *Dictyostelium discoideum* | 44689 | protists | ddiscoideum_eg_gene | G; U |
| *Drosophila melanogaster* | 7227 | ensembl | dmelanogaster_gene_ensembl | G; U |
| *Drosophila pseudoobscura* | 7237 | metazoa | dpgca009870125v2rs_eg_gene | G; U |
| *Equus caballus* | 9796 | ensembl | ecaballus_gene_ensembl | G; U |
| *Gallus gallus* | 9031 | ensembl | ggallus_gene_ensembl | G; U |
| *Glycine max* | 3847 | plants | gmax_eg_gene | G; U |
| *Homo sapiens* | 9606 | ensembl | hsapiens_gene_ensembl | G; U; P |
| *Hordeum vulgare subsp. vulgare* | 4513 | plants | hv10tj18_eg_gene | G; U |
| *Macaca mulatta* | 9544 | ensembl | mmulatta_gene_ensembl | G; U |
| *Monodelphis domestica* | 13616 | ensembl | mdomestica_gene_ensembl | G; U |
| *Mus musculus* | 10090 | ensembl | mmusculus_gene_ensembl | G; U; P |
| *Musa acuminata AAA Group* | 4641 | plants | macuminata_eg_gene | G; U |
| *Oryctolagus cuniculus* | 9986 | ensembl | ocuniculus_gene_ensembl | G; U |
| *Oryza sativa Japonica Group* | 39947 | plants | osativa_eg_gene | G; U |
| *Oryza sativa* | 4530 | plants | osarc_eg_gene | G; U |
| *Ovis aries* | 9940 | ensembl | oaries_gene_ensembl | G; U |
| *Papio anubis* | 9555 | ensembl | panubis_gene_ensembl | G; U |
| *Populus trichocarpa* | 3694 | plants | ptrichocarpa_eg_gene | G; U |
| *Rattus norvegicus* | 10116 | ensembl | rnorvegicus_gene_ensembl | G; U; P |
| *Saccharomyces cerevisiae* | 4932 | ensembl | scerevisiae_gene_ensembl | G; U |
| *Schistosoma mansoni* | 6183 | metazoa | smansoni_eg_gene | G; U |
| *Setaria italica* | 4555 | plants | sitalica_eg_gene | G; U |
| *Solanum lycopersicum* | 4081 | plants | slgca000188115v5cm_eg_gene | G; U |
| *Solanum tuberosum* | 4113 | plants | stuberosum_eg_gene | G; U |
| *Sorghum bicolor* | 4558 | plants | sbicolor_eg_gene | G; U |
| *Sus scrofa* | 9823 | ensembl | sscrofa_gene_ensembl | G; U; P |
| *Theobroma cacao* | 3641 | plants | tcacao_eg_gene | G; U |
| *Trifolium pratense* | 57577 | plants | tpratense_eg_gene | G; U |
| *Triticum aestivum* | 4565 | plants | taestivum_eg_gene | G; U |
| *Vitis vinifera* | 29760 | plants | vvinifera_eg_gene | G; U |
| *Xenopus tropicalis* | 8364 | ensembl | xtropicalis_gene_ensembl | G; U |
| *Zea mays* | 4577 | plants | zmays_eg_gene | G; U |
| *Zea mays subsp. mays* | 4577 | plants | zmays_eg_gene | G; U |

**Table S2.** *Arabidopsis thaliana* reference genes for the Calvin-Benson cycle (CBC) and tricarboxylic acid (TCA) cycle.

| Gene ID | Gene name | Description | Amino acid residues | Reaction ID | Reaction EC | Enzymatic activity | Evidence (AraCyc) | MS count |
|---|---|---|---|---|---|---|---|---|
| AT5G38430 | RBCS1B | Ribulose bisphosphate carboxylase (small chain) family protein | 181 | RIBULOSE-BISPHOSPHATE-CARBOXYLASE-RXN | 4.1.1.39 | ribulose-bisphosphate carboxylase | EV-COMP | 26x |
| AT5G38420 | RBCS2B | Ribulose bisphosphate carboxylase (small chain) family protein | 181 | RIBULOSE-BISPHOSPHATE-CARBOXYLASE-RXN | 4.1.1.39 | ribulose-bisphosphate carboxylase | EV-COMP | 22x |
| AT5G38410 | RBCS3B | Ribulose bisphosphate carboxylase (small chain) family protein | 181 | RIBULOSE-BISPHOSPHATE-CARBOXYLASE-RXN | 4.1.1.39 | ribulose-bisphosphate carboxylase | EV-COMP | 20x |
| AT1G67090 | RBCS1A | ribulose bisphosphate carboxylase small chain 1A | 180 | RIBULOSE-BISPHOSPHATE-CARBOXYLASE-RXN | 4.1.1.39 | ribulose-bisphosphate carboxylase | EV-COMP | 40x |
| AT1G56190 | PGK2 | Phosphoglycerate kinase family protein | 478 | PHOSGLYPHOS-RXN | 2.7.2.3 | phosphoglycerate kinase | EV-COMP | 22x |
| AT3G12780 | PGK1 | phosphoglycerate kinase 1 | 481 | PHOSGLYPHOS-RXN | 2.7.2.3 | phosphoglycerate kinase | EV-COMP | 28x |
| AT1G42970 | GAPB | glyceraldehyde-3-phosphate dehydrogenase B subunit | 447 | 1.2.1.13-RXN | 1.2.1.13 | glyceraldehyde-3-phosphate dehydrogenase (NADP+) (phosphorylating) | EV-AS | 33x |
| AT3G26650 | GAPA | glyceraldehyde 3-phosphate dehydrogenase A subunit | 396 | 1.2.1.13-RXN | 1.2.1.13 | glyceraldehyde-3-phosphate dehydrogenase (NADP+) (phosphorylating) | EV-AS | 29x |
| AT1G12900 | GAPA-2 | glyceraldehyde 3-phosphate dehydrogenase A subunit 2 | 399 | 1.2.1.13-RXN | 1.2.1.13 | glyceraldehyde-3-phosphate dehydrogenase (NADP+) (phosphorylating) | EV-AS | 27x |
| AT2G21170 | pdTPI | triosephosphate isomerase | 315 | TRIOSEPISOMERIZATION-RXN | 5.3.1.1 | triose-phosphate isomerase | EV-EXP | 20x |
| AT3G55800 | SBPASE | sedoheptulose-bisphosphatase | 393 | SEDOHEPTULOSE-BISPHOSPHATASE-RXN | 3.1.3.37 | sedoheptulose-bisphosphatase | EV-EXP | 30x |
| AT2G45290 | TKL2 | Transketolase | 741 | 1TRANSKETO-RXN | 2.2.1.1 | transketolase | EV-EXP | 13x |
| AT3G60750 | TKL1 | Transketolase | 741 | 1TRANSKETO-RXN | 2.2.1.1 | transketolase | EV-EXP | 25x |
| AT3G04790 | EMB3119 | Ribose 5-phosphate isomerase, type A protein | 276 | RIB5PISOM-RXN | 5.3.1.6 | ribose-5-phosphate isomerase | EV-COMP | 27x |
| AT5G44520 | RpiA | NagB/RpiA/CoA transferase-like superfamily protein | 296 | RIB5PISOM-RXN | 5.3.1.6 | ribose-5-phosphate isomerase | EV-COMP | 4x |
| AT2G01140 | PDE345 | Aldolase superfamily protein | 391 | F16ALDOLASE-RXN | 4.1.2.13 | fructose-bisphosphate aldolase | EV-COMP | 27x |
| AT2G21330 | FBA1 | fructose-bisphosphate aldolase 1 | 399 | F16ALDOLASE-RXN | 4.1.2.13 | fructose-bisphosphate aldolase | EV-COMP | 37x |
| AT4G38970 | FBA2 | fructose-bisphosphate aldolase 2 | 398 | F16ALDOLASE-RXN | 4.1.2.13 | fructose-bisphosphate aldolase | EV-COMP | 35x |
| AT4G26530 | FBA5 | Aldolase superfamily protein | 358 | F16ALDOLASE-RXN | 4.1.2.13 | fructose-bisphosphate aldolase | EV-AS | 9x |
| AT5G64380 | AT5G64380 | Inositol monophosphatase family protein | 404 | F16BDEPHOS-RXN | 3.1.3.11 | fructose-bisphosphatase | EV-COMP | 5x |
| AT3G54050 | HCEF1 | high cyclic electron flow 1 | 417 | F16BDEPHOS-RXN | 3.1.3.11 | fructose-bisphosphatase | EV-IC | 24x |
| AT5G61410 | RPE | D-ribulose-5-phosphate-3-epimerase | 281 | RIBULP3EPIM-RXN | 5.1.3.1 | ribulose-phosphate 3-epimerase | EV-COMP | 20x |
| AT1G32060 | PRK | phosphoribulokinase | 395 | PHOSPHORIBULOKINASE-RXN | 2.7.1.19 | phosphoribulokinase | EV-EXP | 31x |
| AT2G47510 | FUM1 | fumarase 1 | 492 | FUMHYDR-RXN | 4.2.1.2 | fumarate hydratase | EV-COMP | 19x |
| AT1G53240 | mMDH1 | Lactate/malate dehydrogenase family protein | 341 | MALATE-DEH-RXN | 1.1.1.37 | malate dehydrogenase | EV-EXP | 23x |
| AT3G15020 | mMDH2 | Lactate/malate dehydrogenase family protein | 341 | MALATE-DEH-RXN | 1.1.1.37 | malate dehydrogenase | EV-EXP | 15x |
| AT2G13560 | NAD-ME1 | NAD-dependent malic enzyme 1 | 623 | MALATE-DEH-RXN | 1.1.1.37 | malate dehydrogenase (oxaloacetate-decarboxylating) | EV-EXP | 14x |
| AT4G00570 | NAD-ME2 | NAD-dependent malic enzyme 2 | 607 | MALATE-DEH-RXN | 1.1.1.37 | malate dehydrogenase (oxaloacetate-decarboxylating) | EV-EXP | 17x |
| AT3G60100 | CSY5 | citrate synthase 5 | 464 | CITSYN-RXN | 2.3.3.1/2.3.3.3/2.3.3.16 | citrate (Re)-synthase | EV-COMP | 6x |
| AT2G44350 | CSY4 | Citrate synthase family protein | 473 | CITSYN-RXN | 2.3.3.1/2.3.3.3/2.3.3.16 | citrate (Re)-synthase | EV-EXP | 20x |
| AT4G26970 | ACO2 | aconitase 2 | 995 | ACONITATEDEHYDR-RXN | No EC# | citrate hydro-lyase (cis-aconitate-forming) | EV-EXP | 17x |
| AT2G05710 | ACO3 | aconitase 3 | 990 | ACONITATEDEHYDR-RXN | No EC# | citrate hydro-lyase (cis-aconitate-forming) | EV-EXP | 21x |
| AT5G54950 | ACO-like | Aconitase family protein | 74 | ACONITATEDEHYDR-RXN | No EC# | aconitate hydratase | EV-COMP | 2x |
| AT2G17130 | IDH-II | isocitrate dehydrogenase subunit 2 | 367 | ISOCITRATE-DEHYDROGENASE-NAD+-RXN | 1.1.1.41/1.1.1.286 | isocitrate--homoisocitrate dehydrogenase | EV-EXP | 13x |
| AT5G03290 | IDH-V | isocitrate dehydrogenase V | 374 | ISOCITRATE-DEHYDROGENASE-NAD+-RXN | 1.1.1.41/1.1.1.286 | isocitrate--homoisocitrate dehydrogenase | EV-EXP | 19x |
| AT4G35650 | IDH-III | isocitrate dehydrogenase III | 368 | ISOCITRATE-DEHYDROGENASE-NAD+-RXN | 1.1.1.41/1.1.1.286 | isocitrate--homoisocitrate dehydrogenase | EV-EXP | 5x |
| AT4G35260 | IDH-I | isocitrate dehydrogenase 1 | 367 | ISOCITRATE-DEHYDROGENASE-NAD+-RXN | 1.1.1.41/1.1.1.286 | isocitrate--homoisocitrate dehydrogenase | EV-EXP | 18x |
| AT3G09810 | IDH-VI | isocitrate dehydrogenase VI | 374 | ISOCITRATE-DEHYDROGENASE-NAD+-RXN | 1.1.1.41/1.1.1.286 | isocitrate--homoisocitrate dehydrogenase | EV-EXP | 16x |
| AT2G20420 | AT2G20420 | ATP citrate lyase (ACL) family protein | 421 | SUCCCOASYN-RXN | 6.2.1.5 | succinate--CoA ligase (ADP-forming) | EV-COMP | 24x |
| AT5G08300 | AT5G08300 | Succinyl-CoA ligase, alpha subunit | 347 | SUCCCOASYN-RXN | 6.2.1.5 | succinate--CoA ligase (ADP-forming) | EV-COMP | 20x |
| AT5G23250 | AT5G23250 | Succinyl-CoA ligase, alpha subunit | 341 | SUCCCOASYN-RXN | 6.2.1.5 | succinate--CoA ligase (ADP-forming) | EV-COMP | 14x |
| AT5G09600 | SDH3-1 | succinate dehydrogenase 3-1 | 213 | RXN-14971 | 1.3.5.1 | succinate dehydrogenase | EV-COMP | 6x |
| AT3G27380 | SDH2-1 | succinate dehydrogenase 2-1 | 279 | RXN-14971 | 1.3.5.1 | succinate dehydrogenase | EV-COMP | 16x |
| AT2G18450 | SDH1-2 | succinate dehydrogenase 1-2 | 632 | RXN-14971 | 1.3.5.1 | succinate dehydrogenase | EV-COMP | 8x |
| AT5G62575 | SDH7B | unknown | 99 | RXN-14971 | 1.3.5.1 | succinate dehydrogenase | EV-COMP | 10x |
| AT2G46390 | SDH8 | succinate dehydrogenase subunit 8 | 46 | RXN-14971 | 1.3.5.1 | succinate dehydrogenase | EV-EXP | 2x |
| AT3G47833 | SDH7A | succinate dehydrogenase subunit 7A | 93 | RXN-14971 | 1.3.5.1 | succinate dehydrogenase | EV-EXP | 11x |
| AT1G08480 | SDH6 | succinate dehydrogenase subunit 6 | 142 | RXN-14971 | 1.3.5.1 | succinate dehydrogenase | EV-EXP | 17x |
| AT1G47420 | SDH5 | succinate dehydrogenase subunit 5 | 257 | RXN-14971 | 1.3.5.1 | succinate dehydrogenase | EV-EXP | 24x |
| AT2G46505 | SDH4 | succinate dehydrogenase subunit 4 | 151 | RXN-14971 | 1.3.5.1 | succinate dehydrogenase | EV-EXP | 8x |
| AT4G32210 | SDH3-2 | succinate dehydrogenase 3-2 | 213 | RXN-14971 | 1.3.5.1 | succinate dehydrogenase | EV-EXP | 4x |
| AT5G40650 | SDH2-2 | succinate dehydrogenase 2-2 | 280 | RXN-14971 | 1.3.5.1 | succinate dehydrogenase | EV-EXP | 14x |
| AT5G66760 | SDH1-1 | succinate dehydrogenase 1-1 | 634 | RXN-14971 | 1.3.5.1 | succinate dehydrogenase | EV-EXP | 25x |

**Table S3.** Maize homologs of Arabidopsis CBC and TCA cycle query genes identified by iGEX. No hits were found for the following accession IDs: AT4G26970, AT5G54950, AT4G35650, AT5G09600, AT4G32210, AT5G62575, AT2G46390, AT3G47833, AT1G08480, AT1G47420, AT2G46505.

| Query locus | Zea mays accession | Zea mays gene name | Query coverage (%) | Confidence | iGEX-assigned name |
|---|---|---|---|---|---|
| AT5G38430 | ZM00001EB092540 | RBCS_1 | 62.9 | medium | Zm-CBC_RBCS1B.1 |
| AT5G38430 | ZM00001EB197410 | RBCS | 62.4 | medium | Zm-CBC_RBCS1B.2 |
| AT1G56190 | ZM00001EB352550 | | 80.3 | medium | Zm-CBC_PGK2.1 |
| AT1G56190 | ZM00001EB291670 | | 79.9 | medium | Zm-CBC_PGK2.2 |
| AT1G56190 | ZM00001EB150720 | | 69.7 | medium | Zm-CBC_PGK2.3 |
| AT1G42970 | ZM00001EB002270 | GAPB | 80.1 | medium | Zm-CBC_GAPB |
| AT3G26650 | ZM00001EB424110 | | 84 | medium | Zm-CBC_GAPA.1 |
| AT3G26650 | ZM00001EB424190 | | 79.8 | medium | Zm-CBC_GAPA.2 |
| AT3G26650 | ZM00001EB424150 | | 79.5 | medium | Zm-CBC_GAPA.3 |
| AT3G26650 | ZM00001EB080840 | | 47.5 | medium | Zm-CBC_GAPA.4 |
| AT2G21170 | ZM00001EB103580 | TPIC_4 | 71.8 | high | Zm-CBC_pdTPI.1 |
| AT2G21170 | ZM00001EB319770 | | 71.4 | high | Zm-CBC_pdTPI.2 |
| AT3G55800 | ZM00001EB147750 | shbp1 | 75.8 | medium | Zm-CBC_SBPASE |
| AT2G45290 | ZM00001EB389880 | | 71 | medium | Zm-CBC_TKL2 |
| AT3G04790 | ZM00001EB301210 | | 67.5 | high | Zm-CBC_EMB3119 |
| AT5G44520 | ZM00001EB213570 | | 46 | high | Zm-CBC_RpiA |
| AT2G01140 | ZM00001EB126080 | | 81.9 | medium | Zm-CBC_PDE345.1 |
| AT2G01140 | ZM00001EB334740 | | 76.5 | medium | Zm-CBC_PDE345.2 |
| AT2G01140 | ZM00001EB010390 | | 76 | medium | Zm-CBC_PDE345.3 |
| AT2G01140 | ZM00001EB085480 | | 76 | medium | Zm-CBC_PDE345.4 |
| AT2G01140 | ZM00001EB188850 | | 74.8 | medium | Zm-CBC_PDE345.5 |
| AT2G21330 | ZM00001EB408010 | | 82.5 | medium | Zm-CBC_FBA1.1 |
| AT2G21330 | ZM00001EB200910 | | 80.7 | medium | Zm-CBC_FBA1.2 |
| AT4G26530 | ZM00001EB386660 | | 79.6 | medium | Zm-CBC_FBA5 |
| AT5G64380 | ZM00001EB389720 | F16P1 | 64.3 | high | Zm-CBC_AT5G64380 |
| AT3G54050 | ZM00001EB011940 | | 73.8 | medium | Zm-CBC_HCEF1 |
| AT5G61410 | ZM00001EB005330 | | 76.2 | medium | Zm-CBC_RPE.1 |
| AT5G61410 | ZM00001EB402390 | | 75.4 | medium | Zm-CBC_RPE.2 |
| AT1G32060 | ZM00001EB251520 | KPPR_0 | 82.5 | medium | Zm-CBC_PRK.1 |
| AT1G32060 | ZM00001EB188120 | | 81.5 | medium | Zm-CBC_PRK.2 |
| AT1G32060 | ZM00001EB072030 | | 74.8 | medium | Zm-CBC_PRK.3 |
| AT2G47510 | ZM00001EB016240 | FUM2 | 83.3 | medium | Zm-TCA_FUM1 |
| AT1G53240 | ZM00001EB158270 | MMDHI | 82.1 | medium | Zm-TCA_mMDH1.1 |
| AT1G53240 | ZM00001EB344340 | | 79.8 | medium | Zm-TCA_mMDH1.2 |
| AT1G53240 | ZM00001EB296550 | | 79.1 | medium | Zm-TCA_mMDH1.3 |
| AT2G13560 | ZM00001EB321980 | | 78.7 | medium | Zm-TCA_NAD-ME1 |
| AT4G00570 | ZM00001EB220780 | | 74.5 | medium | Zm-TCA_NAD-ME2 |
| AT3G60100 | ZM00001EB206100 | | 78.5 | medium | Zm-TCA_CSY5.1 |
| AT3G60100 | ZM00001EB236450 | | 78.5 | medium | Zm-TCA_CSY5.2 |
| AT3G60100 | ZM00001EB206090 | | 74 | medium | Zm-TCA_CSY5.3 |
| AT2G05710 | ZM00001EB394900 | | 31.8 | medium | Zm-TCA_ACO3 |
| AT2G17130 | ZM00001EB183760 | | 82.3 | medium | Zm-TCA_IDH-II.1 |
| AT2G17130 | ZM00001EB246080 | | 81.7 | medium | Zm-TCA_IDH-II.2 |
| AT2G17130 | ZM00001EB424940 | | 80.9 | medium | Zm-TCA_IDH-II.3 |
| AT5G03290 | ZM00001EB013280 | | 92.5 | medium | Zm-TCA_IDH-V.1 |
| AT5G03290 | ZM00001EB128830 | | 81.1 | medium | Zm-TCA_IDH-V.2 |
| AT5G03290 | ZM00001EB332780 | | 80.5 | medium | Zm-TCA_IDH-V.3 |
| AT5G03290 | ZM00001EB013290 | | 75.5 | medium | Zm-TCA_IDH-V.4 |
| AT2G20420 | ZM00001EB247500 | | 84.6 | medium | Zm-TCA_AT2G20420.1 |
| AT2G20420 | ZM00001EB185030 | | 84.1 | medium | Zm-TCA_AT2G20420.2 |
| AT2G20420 | ZM00001EB277730 | | 74.6 | medium | Zm-TCA_AT2G20420.3 |
| AT2G20420 | ZM00001EB283930 | | 68.1 | medium | Zm-TCA_AT2G20420.4 |
| AT5G08300 | ZM00001EB325420 | | 80.7 | medium | Zm-TCA_AT5G08300.1 |
| AT5G08300 | ZM00001EB107590 | Os07g0577700_1 | 60.6 | medium | Zm-TCA_AT5G08300.2 |
| AT3G27380 | ZM00001EB097190 | | 70.9 | medium | Zm-TCA_SDH2-1.1 |
| AT3G27380 | ZM00001EB358070 | | 70.2 | medium | Zm-TCA_SDH2-1.2 |
| AT3G27380 | ZM00001EB175570 | | 65.7 | medium | Zm-TCA_SDH2-1.3 |
| AT3G27380 | ZM00001EB261270 | SDH2-1_3 | 55.6 | medium | Zm-TCA_SDH2-1.4 |
| AT3G27380 | ZM00001EB173490 | sdh2-rps14 | 54.8 | medium | Zm-TCA_SDH2-1.5 |
| AT2G18450 | ZM00001EB117970 | SDH1_0 | 87.6 | medium | Zm-TCA_SDH1-2.1 |
| AT2G18450 | ZM00001EB299550 | | 87.2 | medium | Zm-TCA_SDH1-2.2 |
| AT2G18450 | ZM00001EB337870 | | 72.3 | medium | Zm-TCA_SDH1-2.3 |
| AT2G18450 | ZM00001EB306180 | | 65.3 | medium | Zm-TCA_SDH1-2.4 |
